# Model of leaf biomass partitioning coefficient in different leaf ranks of rapeseed (*Brassica napus* L.) main stem

**DOI:** 10.1101/2025.07.27.667074

**Authors:** Weixin Zhang, Wenyu Zhang, Qian Wu, Chuanliang Sun, Daokuo Ge, Jing Cao, Hong Li, Hongxin Cao

## Abstract

Leaf growth is a dynamic process, and understanding the number of main-stem leaves and the distribution of leaf biomass at different development stages can provide a theoretical basis for the development of functional-structural rapeseed model. To address the gap in research on the leaf biomass partitioning coefficient model in the crop growth model, we developed the leaf biomass partitioning coefficient model for the main-stem at different leaf ranks in rapeseed. The outdoor experiment with cultivars, Ningyou 18 (V1, conventional), Ningyou 16 (V2, conventional), and Ningza 19 (V3, hybrid), and designed treatment of cultivars × fertilizer × transplanting density in the 2012-2015 growth season. The model was fitted by calculating the ratio of leaf biomass at different leaf ranks on the main-stem to the total leaf biomass on the main-stem, and normalizing the leaf ranks into (0-1] interval. We parameterized the model variables to elucidate how cultivar and environmental conditions influence the leaf biomass partitioning coefficients for different leaf ranks. The descriptive models were validated with independent experimental data, the correlation (*r*) of simulation and observation values all greater than 0.9 with significant levels at *p* < 0.001. The absolute values of the average absolute difference (*d*_*a*_) are all less than 0.011 g g^−1^. The ratio of *d*_*a*_ to the average observation (*d*_*ap*_) for the normalized leaf rank, and leaf biomass partitioning coefficient at seedling stage, and budding and bolting stage, and blooming stage, and mature stage are 13.599%, 11.236%, 11.312%, 3.077%, and 8.661%, respectively. The *RMSE* values are all less than 0.032 g g^−1^. The models developed in this paper have good performance and reliability for predicting leaf biomass partitioning coefficient of main-stem at different leaf ranks in rapeseed. This study integrates the leaf morphology model of rapeseed with the rapeseed growth model through biomass partitioning, contributing to the development of the functional-structural rapeseed model.

## 1. Introduction

As principal photosynthetic organs, leaves perform critical physiological processes including light energy conversion, gaseous exchange, and photoassimilate production through coordinated photosynthetic and respiratory activities (Niinemets, 2023). The ontogeny of leaves initiates through leaf primordia formation, a developmental process exhibiting phyllotactic coordination with branch, floral bud, and pod organogenesis (Luo *et al*., 2018). Empirical evidence demonstrates that photosynthetic enhancement induces carbohydrate surplus, creating positive feedback mechanisms that stimulate leaf morphogenesis (Raines, 2006). In *Brassica napus*, the architectural configuration of plants is fundamentally determined by leaf morphological characteristics, phyllotaxis patterns, and population density. Strategic optimization of foliar quantity and biomass distribution patterns presents opportunities for improving canopy architecture, augmenting photosynthetic performance, rationalizing biomass allocation, and ultimately enhancing crop productivity and quality.

Contemporary understanding of biomass allocation mechanisms reveals that assimilate partitioning among plant organs critically influences growth dynamics and yield formation (Marcelis and Heuvelink, 2007). Traditional allometric models, despite their widespread adoption in crop simulation platforms, employ static organ growth ratios that constrain predictive accuracy to specific environmental regimes (Wilson, 1988). While providing reasonable approximations under controlled conditions, these empirically-derived approaches inadequately capture the dynamic reallocation patterns observed under fluctuating resource availability (Marcelis, 1993b). More critically, their inherent empirical nature fails to explain emerging evidence of carbon-nitrogen stoichiometric regulation in biomass partitioning (Van der Werf *et al*., 1993; Thornley and Johnson, 1990). Consequently, quantitative and mechanistic insights into biomass allocation are often limited. While these models may closely align with experimental results, they often describe biomass distribution within a narrow range of growth conditions and fail to account for dynamic fluctuations in biomass allocation (Wilson, 1988).

This conceptual limitation has driven the development of hybrid modeling paradigms. Canonical modeling approaches attempt to bridge empirical and mechanistic extremes by incorporating semi-quantitative physiological principles (Renton *et al*., 2005), enabling simulation of complex allocation dynamics without exhaustive parameterization. However, validation studies reveal that sink strength modulation-defined as the organ-specific growth potential under non-limiting assimilate supply-exerts predominant control over partitioning patterns (Marcelis, 1996). Quantitative characterization of sink strength-defined as the maximum potential growth rate under non-limiting assimilate supply-provides a physiologically grounded framework for modeling inter-organ allocation dynamics (Drouet and Pagès, 2007; Kang *et al*., 2011). The transport resistance model (Thornley, 1976; Dewar, 1993; Minchin and Lacointe, 2005) offers detailed mechanistic representation of assimilate flux from source to sink tissues. However, implementation challenges arising from parameterization complexity and limited operational impact assessments have restricted its adoption in production-scale crop models.

Notably, a critical knowledge gap persists regarding leaf biomass partitioning along ontogenetic gradients in rapeseed (*Brassica napus* L.). Current models predominantly address organ-level allocation rather than intra-organ gradients, despite evidence that leaf rank position influences both photosynthetic competence and senescence patterns. Precise characterization of main stem leaf biomass distribution represents an essential step toward understanding whole-plant resource optimization strategies, particularly for improving stress resilience and carbon sequestration efficiency in breeding programs. Building upon our previous work in rapeseed organ-level partitioning (Zhang *et al*., 2020), this investigation establishes a novel framework for modeling main stem leaf biomass distribution across developmental ranks. Our approach integrates functional-structural plant model principles with sink strength dynamics, aiming to advance precision agriculture applications through improved canopy architecture optimization and resource use efficiency prediction in rapeseed plant cultivation systems.

## 2. Materials and methods

### 2.1. Materials

We used 3 rapeseed cultivars, “Ningyou 18” (V1, conventional), “Ningyou 16” (V2, conventional), and “Ningza 19” (V3, hybrid), bred by the Institute of Economic Crops Research, Jiangsu Academy of Agricultural Sciences.

### 2.2. Methods

#### 2.2.1. Experiments

Four experiments were conducted at the farm of Jiangsu Academy of Agricultural Sciences, Nanjing, China (32.03 ° N, 118.87 ° E), involving different cultivars, transplanting densities, and fertilizer during the 2012–2015 growing seasons. Nanjing is a rainfed area for rapeseed, with an average rainfall of 1106.5 mm, and a relative humidity of 76%. Field rapeseed does not need artificial irrigation, and there is no water-stress when the drainage system is in good condition. The soil type of the experimental area is a hydraulic anthrosol. Soil test results indicated the following: organic carbon, 31.4 g kg^−1^; total nitrogen, 2.03 g kg^−1^; available phosphorus, 20.3 mg kg^−1^; available potassium, 139.0 mg kg^−1^; and pH 7.31.

Experiment 1, cultivar, fertilizer, and transplanting density experiment (2012−2013): The experiment was deployed in split block design with three replications. Three fertilizer levels (N0 = no fertilizer; N1 = 90 kg N ha ^−1^; N2 = 180 kg N ha^−1^) were the whole-plot treatments while cultivar (C1 and C2) and three transplanting densities (D1 = 6×10 ^4^ plant ha^−1^; D2 = 1.2 × 10^5^ plant ha^−1^; D3 = 1.8 × 10^5^ plant ha^−1^) constituted the sub-plots.

Experiment 2, fertilizer and transplanting density experiment (2013−2014): The experiments were deployed in split block design with three replications. Three N fertilizer levels (N0 = no fertilizer; N1 = 90 kg N ha ^−1^; N2 = 180 kg N ha ^−1^) were the whole-plot treatments while cultivar (C1) and three transplanting densities (D1 = 6×10 ^4^ plant ha^−1^; D2 = 1.2 × 10^5^ plant ha^−1^; D3 = 1.8 × 10^5^ plant ha^−1^) constituted the sub-plots.

Experiment 3, cultivar experiment (2013−2014): The experiment was a randomized complete block design with three cultivars (V1, V2 and V3) and three replications. Nitrogen fertilizer included 90 kg N ha ^−1^ (N1), and the transplanting density was 1.2 × 10^5^ plant ha^−1^ (D2).

Experiment 4, cultivar and the fertilizer experiment (2014−2015): The experiment was deployed in split block design with three replications. Five N fertilizer levels (N0 =no fertilizer; N1 = 90 kg N ha^−1^; N2 = 180 kg N ha^−1^; N3 = 270 kg N ha^−1^; N4 = 360 kg N ha^−1^) were the whole-plot treatments while three cultivars (V1 and V3) constituted the sub-plots. The transplanting density was 1.2 × 10^5^ plant ha^−1^ (D2). The specific experimental design shown in Table 1.

**Table 1.**
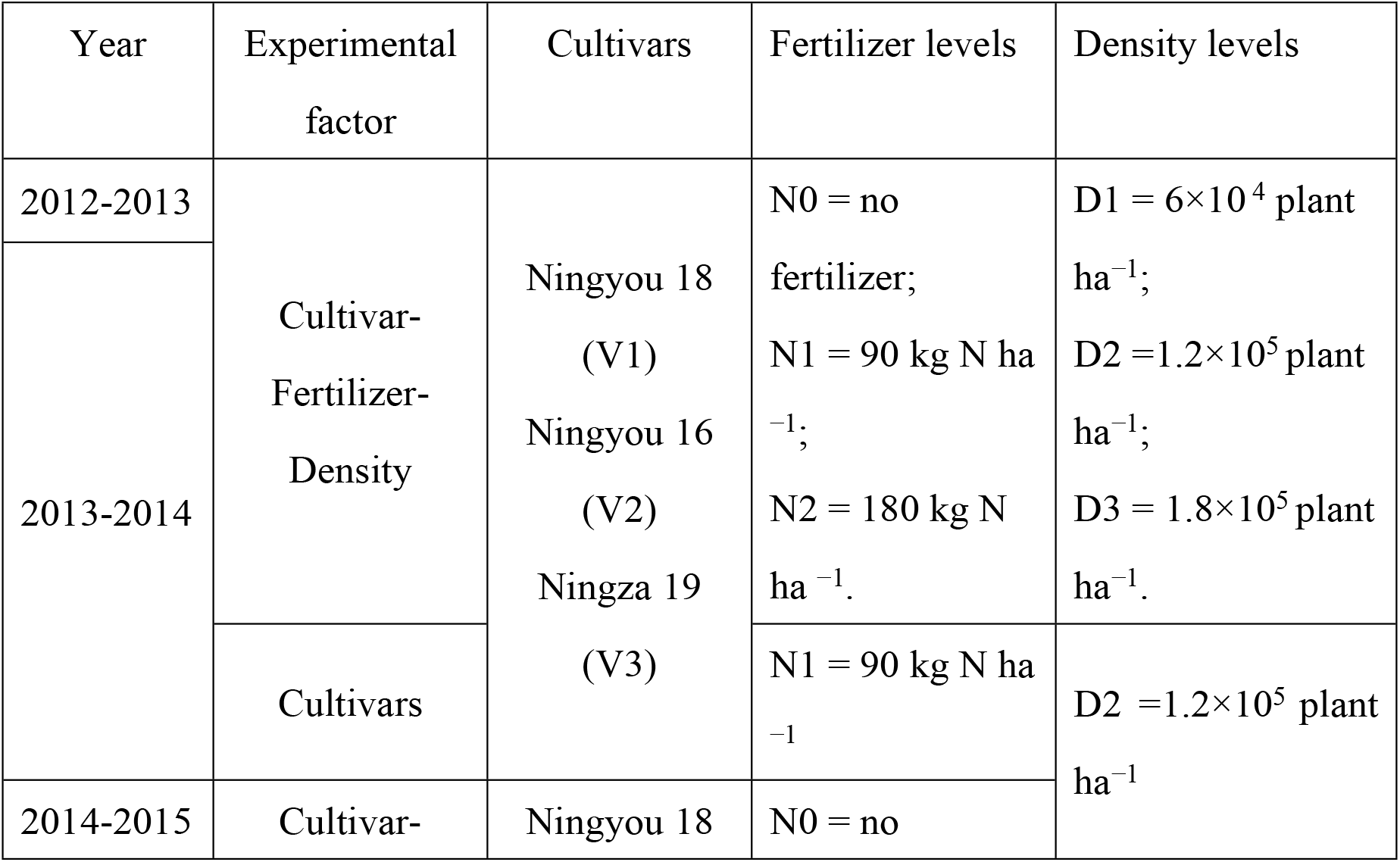
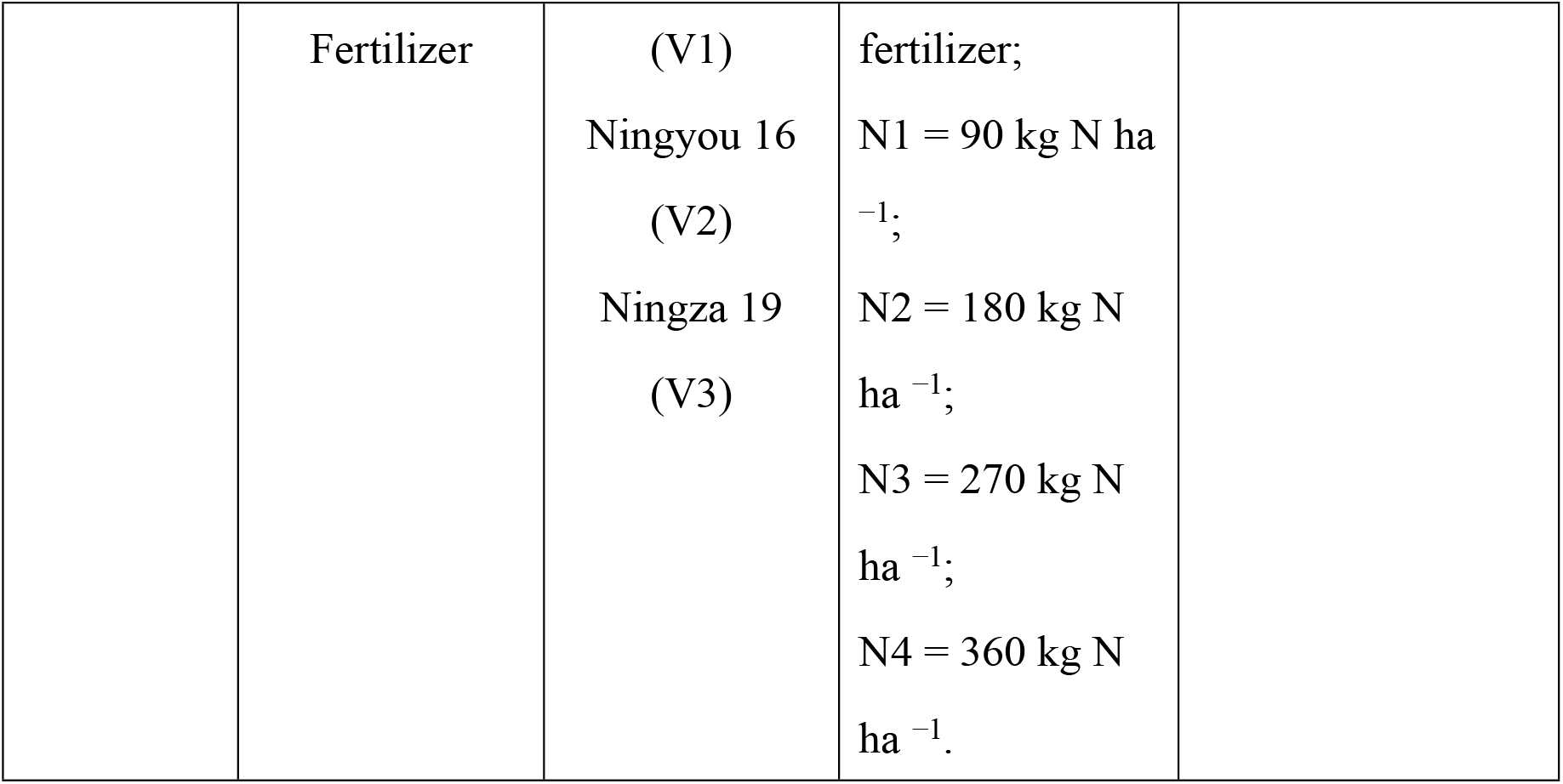
Experimental design.

The plots of all the experiments were arranged randomly with 0.42 m row spacing in 3.99 by 3.5 m area, and the plant spacing was calculated according to row spacing and transplanting density. Fertilizer contained 90 kg P_2_O_5_ ha^−1^ and 90 kg K_2_O ha^−1^ for N1 plots, 180 kg P_2_O_5_ ha^−1^ and 180 kg K_2_O ha^−1^ for N2 plots, 270 kg P_2_O_5_ ha^−1^ and 270 kg K_2_O ha^−1^ for N3 plots, and 360 kg P_2_O_5_ ha^−1^ and 360 kg K_2_O ha^−1^for N4 plots, and 15 kg boron ha^−1^ was used as a foliage spray for N1 to N4 plots after bolting. The ratio of basal to dressing fertilizer at wintering and bolting was 5:3:2. Sowing dates were on 15 Oct. 2011, 8 Oct. 2012, 4 Oct. 2013, and 4 Oct. 2014; and transplanting dates were on 4 Nov. 2011, 9 Nov. 2012, 1 Nov. 2013 and 1 Nov. 2014. The other field management (such as ditching, weeding, pest control, etc.) applied to plots followed the local cultivation practices for high yield in rapeseed.

#### 2.2.2. Sampling and measurements

Before transplanting, the rank of each leaf (i.e. the ordinal position or the leaf order on the stem) on the main-stem of 50 randomly selected seedlings for each plot was marked using a red number stamp. We destructively sampled at least once during each growth stage and chose three to five plants from each of the treatments.

The plant height and the height of each leaf rank were measured. The main-stem leaves were separated by leaf rank, and the other organs including stems, flower buds, flowers, and pods on the main-stem, each primary and secondary branches were separated by leaf rank and organ, respectively. Then, dried in an oven for 30 min at 105 ∘C, then at 80°C until reaching a stable weight, where dry weight was weighed using a 0.001-g electron balance.

#### 2.2.3. Calculation and statistical analysis of relevant indicators

We use physiological day of development (*PDD*) which was used to drive the model (Gao *et al*., 1992), defined as follows:

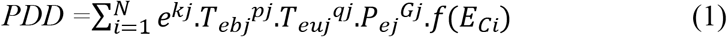

where the N is the number of days covering the stage, the *kj* is basic development parameter which is determined by cultivar heredity, the *T*_*ebj*_ and the *T*_*euj*_ are the effective factors for lower and upper temperature, respectively, the *pj* and the *qj* are the genotypic coefficient of temperature effects, the *P*_*ej*_ is the effective factor of photoperiod, *Gj* is the genotypic coefficient of photoperiod effects, and *f*(*E*_*Ci*_) is the effective function of agronomic practice factors for rapeseed. All the symbols “*j*” above means at the *j*th stage. The calculation of *PDD* in each stage, such as the effects of photoperiod and vernalization, has been briefly described in Appendix A, more details were explained in our existing rapeseed phenology model (Cao *et al*., 2006, 2015; Zhang *et al*., 2020). We normalized *PDD* into [0, 1] interval to calculate and express easily, normalized *PDD* (*nPDD*) 0 means sowing and 1 represents maturity. Biomass partitioning coefficient of leaf (*PC*_*L*_) is defined as follows:

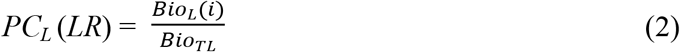

where *Bio*_*L*_ (*i*) is the biomass of the *i*th leaf at the main-stem, and *Bio*_*TL*_ is the total biomass of main-stem leaf.

In order to effectively eliminate the apparent difference of leaf rank and facilitate the construction of the model, we divided the measured leaf rank of each treatment by the total number of leaves to unify it into the (0-1] interval, called normalized leaf rank. The experimental data for the 2013-2014 growing season were used for model development, and the data for the 2012-2013 and 2014-2015 were used for model verification. The multi-way ANOVA was used to analyze the differences in plant height, and the height of the maxi leaf rank, and green leaf and total leaf on the main stem among different experimental treatments SPSS (version 23.0). We arranged the experimental data, fitted grouped data using MS Excel 2016, and find the best fitting formula form using CurveExpert 1.4 using the standard deviation and the correlation coefficient.

#### 2.2.4. Model validation

The models developed in this paper was validated by calculating the correlation (*r*), the root mean square error (*RMSE*), the average absolute difference (*d*_*a*_), and the ratio of da to the average observation (*d*_*ap*_) (Cao *et al*., 2012), and 1:1 line of simulated and observed properties. Some statistical indices are defined as follows:

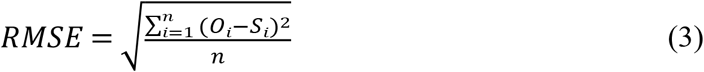

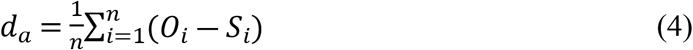

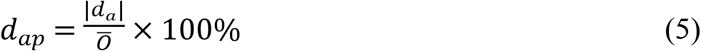

where *i* is sample numbers, *n* is total number of measurements, *n* = *n*-1 when n≥30, *S*_*i*_ is simulated value, and *O*_*i*_ is observed value.

## 3. Results

### 3.1. Variance analysis of leaf biomass partitioning coefficient and green leaf numbers

With the continuous advancement of the growth process of rapeseed plants, each single leaf on the main stem also undergoes a process of elongation, continuation, and senescence. The experimental data showed that the green leaf numbers first increase and then decrease with increasing the normalized *PDD*, and reaching the maximum value when *nPDD* is 0.64, that is, during the early anthesis stage (Fig. 1) under the different treatments.

**Figure 1.**
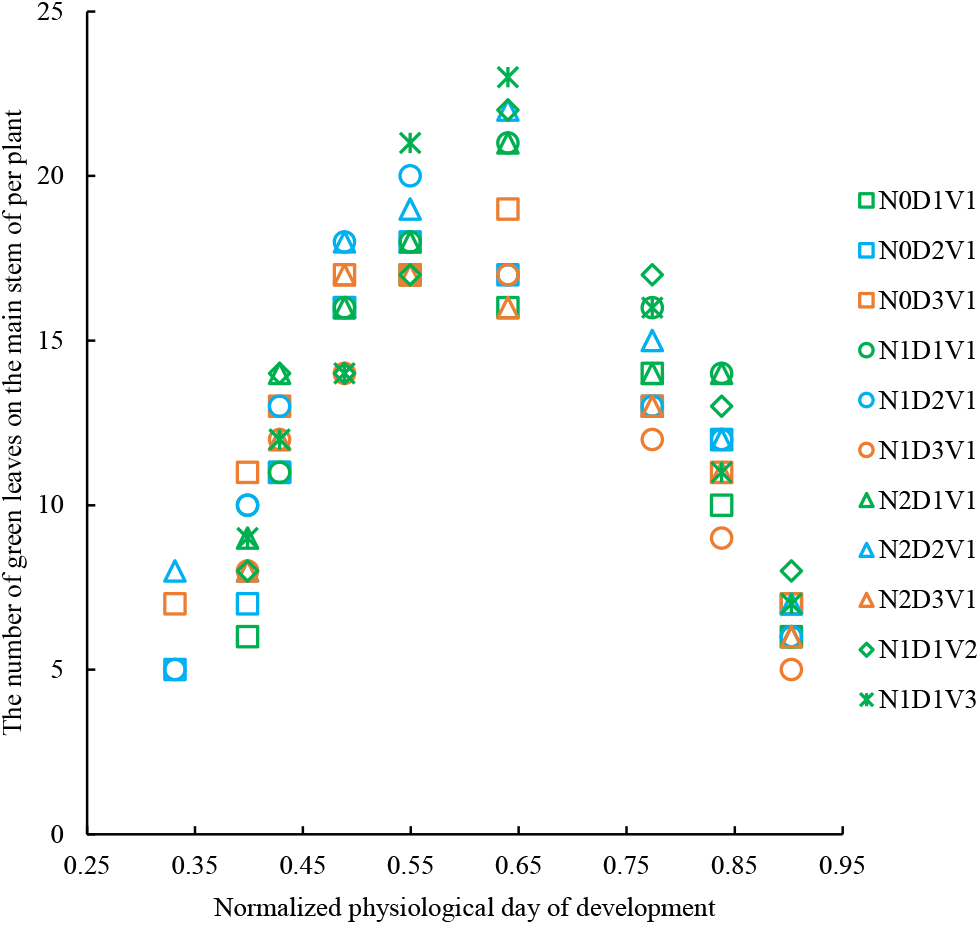
The changes in the green leaf numbers with normalized physiological day of development under different treatments in 2013-2014 growing season.

We analyzed the differences in leaf number and biomass partitioning coefficient under different treatments by using the multi-way ANOVA. The variance analysis result showed that the number of leaves is also affected by cultivar, cultivation measures (fertilization, transplanting density, etc.), and environmental conditions (Table 2).

**Table 2.**
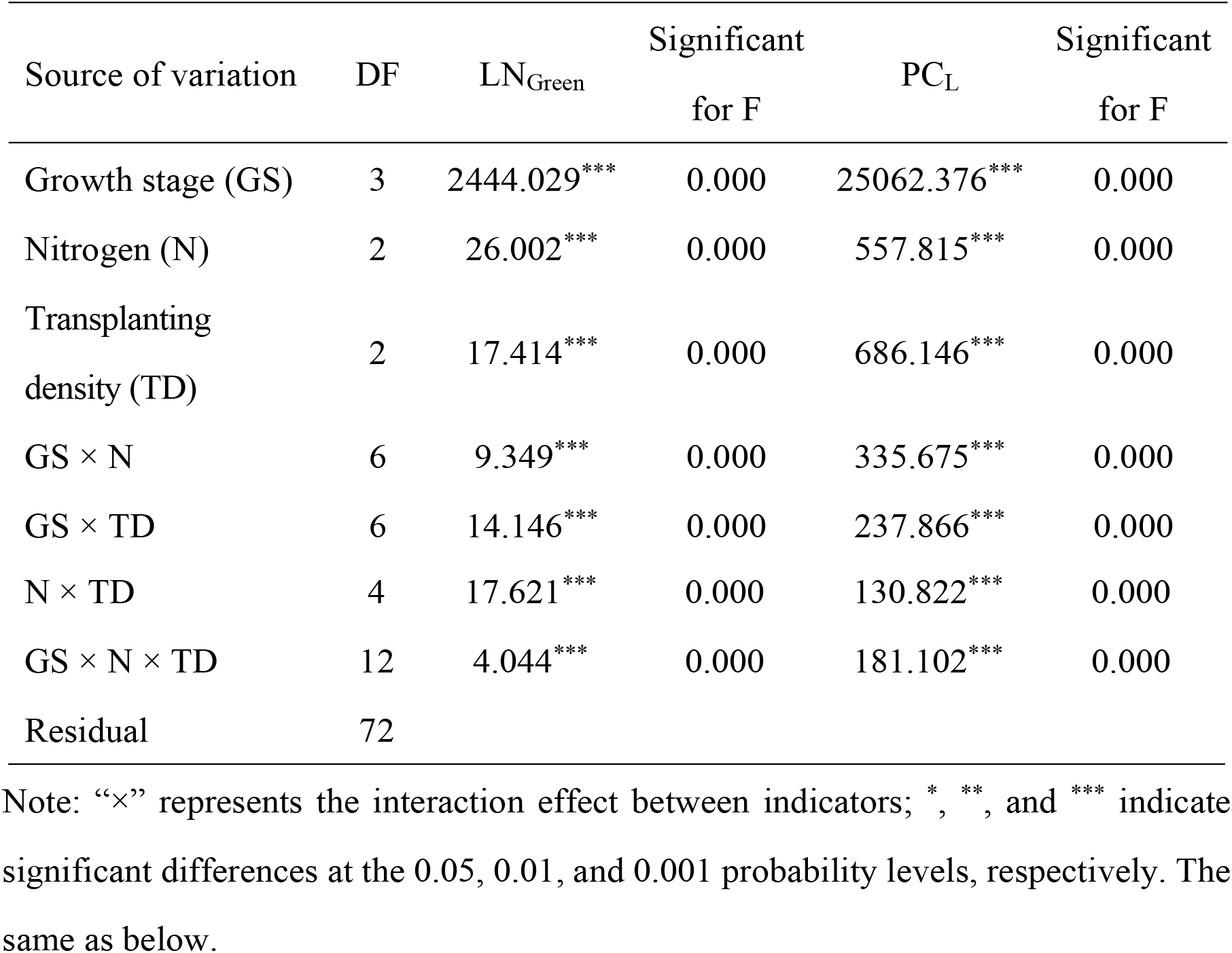
Variance analysis of the effects of growth stage, and cultivar, and fertilizer, and transplanting density, and its interactions on green leaf numbers (LN_Green_), and the leaf biomass partitioning coefficient of rapeseed main stem (PC_L_) at different growth stage. (2013-2014)

### 3.2. The main-stem leaf biomass partitioning coefficient model

#### 3.2.1. The normalized leaf rank model

At any stage of rapeseed growth, there are varying numbers of green leaves on the main stem. However, even for the same cultivar grown in different years, the leaf count can vary, posing challenges for analyzing the change patterns of leaf biomass partitioning coefficients with *PDD*. To address this issue, we introduce the concept of normalized leaf rank (*nLR*) and proceed to investigate the relationship between normalized leaf rank and *PDD* variations. The experimental data showed that the values of *nLR*(*PDD*) of different treatments with normalized *PDD* were fitted by the logarithmic function (Fig. 2, Eq. 6).

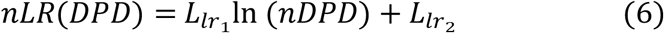

where *nLR*(*DPD*)is the normalized leaf rank at different normalized physiological day of development; *nDPD* is the normalized physiological day of development; 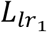, and 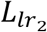 are model parameters. Detailed parameter estimation and significance test are shown in Table 3.

**Table 3.**
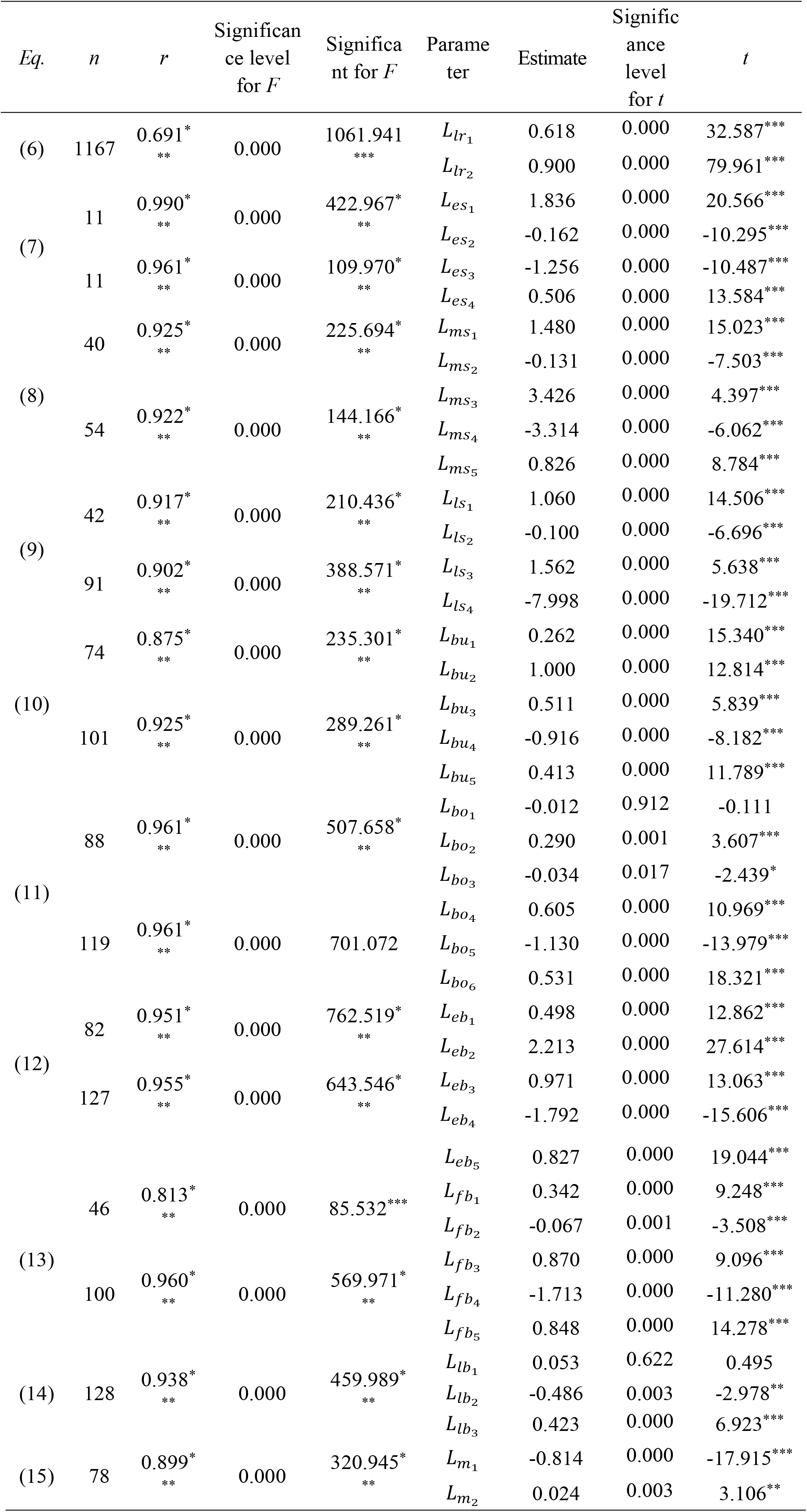
The significant test of the main-stem leaf biomass partitioning coefficient models and their parameters.

**Figure 2.**
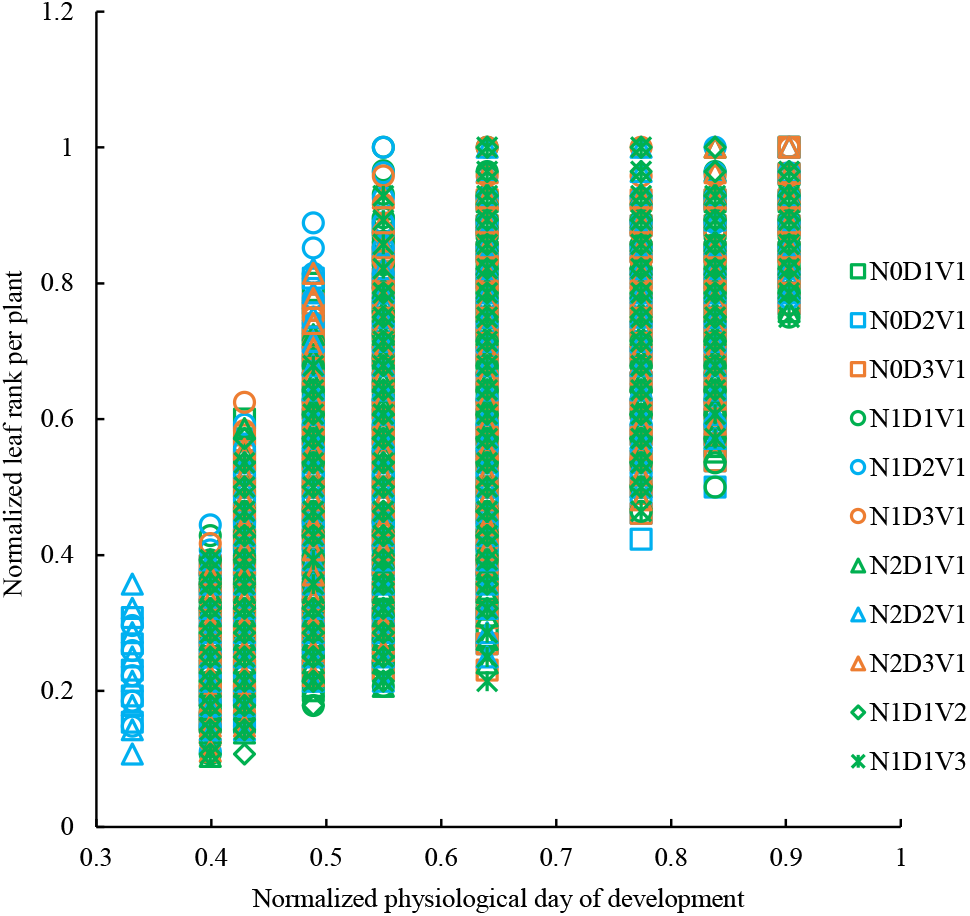
The changes in the normalized leaf rank with normalized physiological day of development under different treatments in 2013-2014 growing season.

#### 3.2.2. Model of leaf biomass partitioning coefficients for the main stem across growth stages

##### 3.2.2.1. Leaf rank boundaries in seedling stage: 0.10-0.63

The seedling stage of winter rapeseed is longer, accounting for almost half or more of the whole growth period. The main-stem leaves are the largest nutrient organ at the seedling stage, and the biomass partitioning coefficient of rapeseed plants is mainly concentrated on the leaves. We divided the seedling stage of rapeseed into pre-winter stage, overwintering stage, and late seedling stage, which can also be called early seedling stage, mid-seedling stage, and late seedling stage. The leaf rank interval in the early seedling stage was (0.10-0.24) and [0.24-0.38), and the leaf rank interval in the mid-seedling stage was (0.10-0.25) and [0.25-0.50], and the late seedling stage was (0.10-0.27) and [0.27-0.63).

The main-stem leaf biomass partitioning coefficient of rapeseed in seedling stage under different treatments all showed a trend of increasing first and then decreasing (Fig. 3). The values of *PC*_*ES*_(*LR*) of different treatments with normalized leaf rank were fitted by the linear function (Equation 7 and Fig. 3A). The values of *PC*_*MS*_(*LR*) of different treatments with normalized leaf rank were fitted by a logarithmic function and a quadratic function (Equation 8 and Fig. 3B). The values of *PC*_*LS*_(*LR*) of different treatments with normalized leaf rank were fitted by a linear function and a power function (Equation 9 and Fig. 3C).

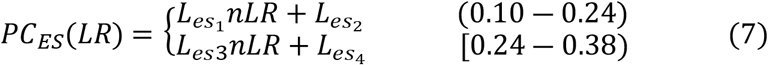

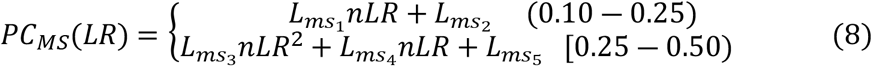

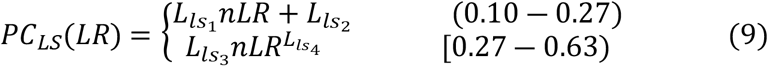

where *PC*_*ES*_(*LR*), *PC*_*MS*_(*LR*), and *PC*_*LS*_(*LR*) are the leaf biomass partitioning coefficient at different leaf rank during the early stage, middle stage, and late stage of rapeseed seedling growth, respectively (g g^−1^); *nLR*(*DPD*) is the normalized leaf rank; 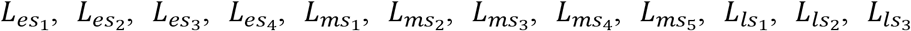 and 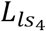 are model parameters. Detailed parameter estimation and significance test are shown in Table 3.

**Figure 3.**
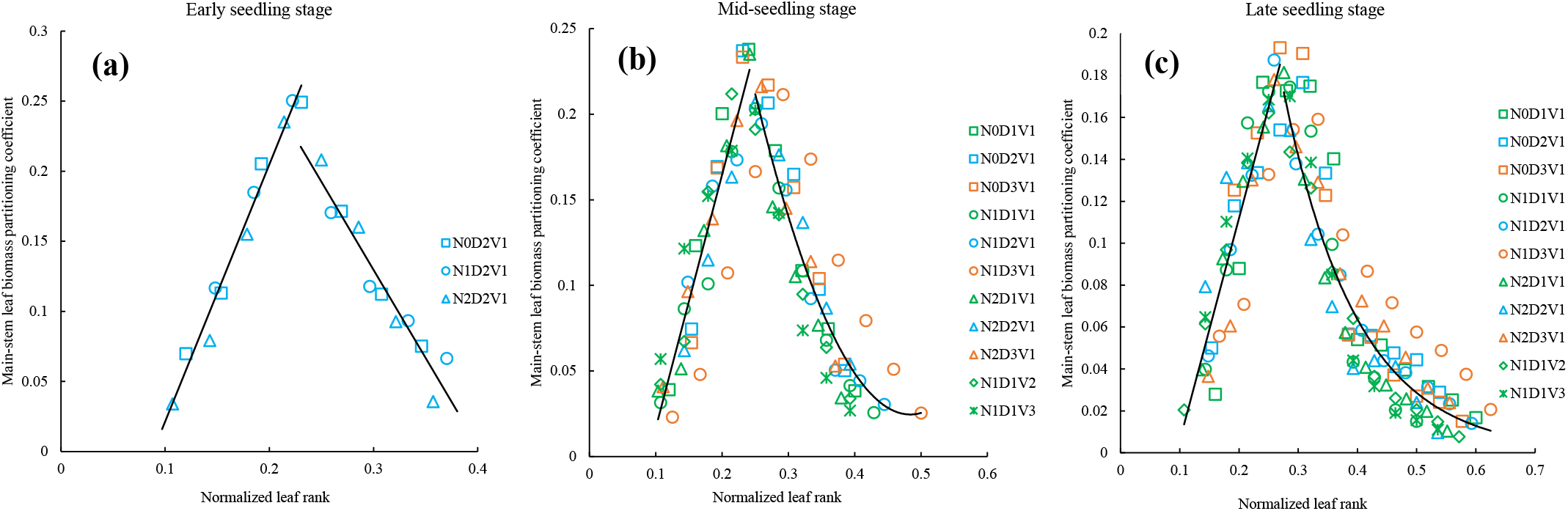
The changes in main-stem leaf biomass partitioning coefficient with normalized leaf rank in the early seedling stage (a) and in the mid-seedling stage (b) and in the late seedling stage (c) under different treatments in 2013-2014 growing season.

##### 3.2.2.2. Leaf rank boundaries in budding and bolting stage: 0.17-1.0

During the budding and bolting stage, there was a rapid accumulation of biomass in rapeseed leaves (Fig. 1A). We divided the budding and bolting stage of rapeseed into budding stage, and bolting stage. The leaf rank interval in the budding stage was (0.17-0.45) and [0.45-0.93), and the leaf rank interval in the bolting stage was (0.21-0.50) and [0.50-1.0].

The main-stem leaf biomass partitioning coefficient of rapeseed in budding and bolting stage under different treatments all showed a trend of increasing first and then decreasing (Fig. 4). The values of PC_BU_(LR) of different treatments with normalized leaf rank were fitted by a linear function and a quadratic function (Equation 10 and Fig. 4A). The values of PC_BO_(LR) of different treatments with normalized leaf rank were both fitted by the quadratic function (Equation 11 and Fig. 4B).

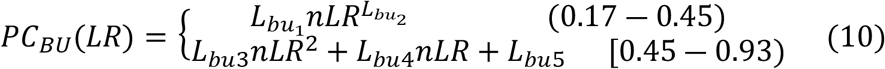

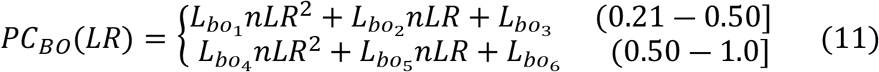

where *PC*_*BU*_(*LR*), and *PC*_*BO*_(*LR*) are the leaf biomass partitioning coefficient at different leaf rank during the budding stage and bolting stage, respectively (g g^−1^); *nLR* is the normalized leaf rank;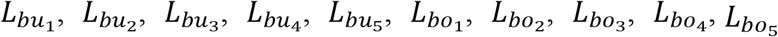, and 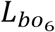 are model parameters. Detailed parameter estimation and significance test are shown in Table 3.

**Figure 4.**
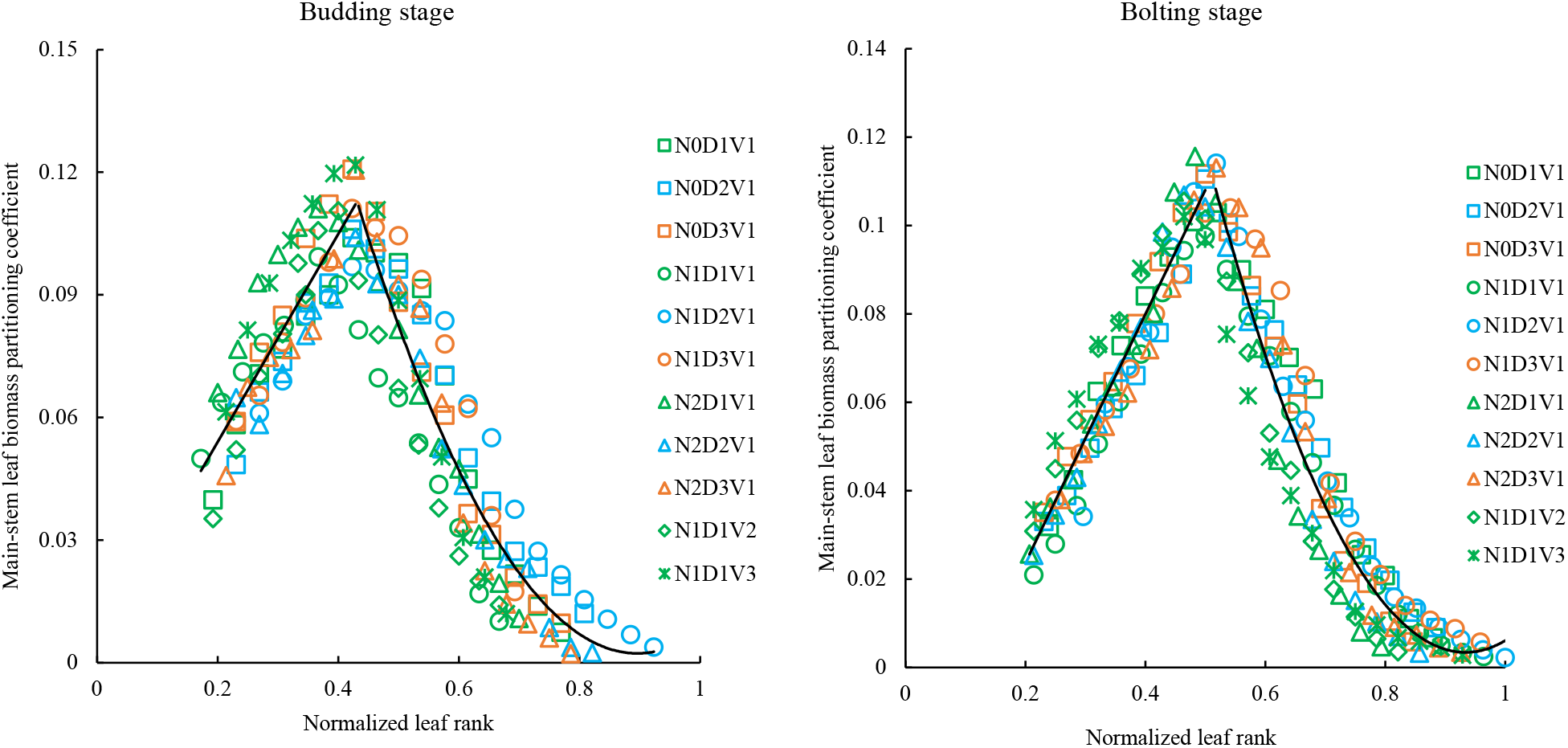
The changes in main-stem leaf biomass partitioning coefficient with normalized leaf rank in the budding and stage (A) and in the bolting stage (B) under different treatments.

##### 3.2.2.3. Leaf rank boundaries in blooming stage: 0.21-1.0

The experimental data showed that the leaf number of rapeseed exhibits a decreasing trend during the blooming stage, consequently leading to a decline in the biomass accumulation of the main stem. Conversely, the leaf biomass partitioning coefficient of rapeseed slightly increased from the early blooming stage to the full blooming stage and then to the late blooming stage as the total leaf number decreased (Fig. 5). We divided the blooming stage of rapeseed into early blooming stage, full blooming stage, and late blooming stage. The leaf rank interval in the early blooming stage was (0.21-0.55) and [0.55-1.0], and the leaf rank interval in the full blooming stage was (0.42-0.60) and [0.60-1.0], and the late blooming stage was [0.50-1.0].

**Figure 5.**
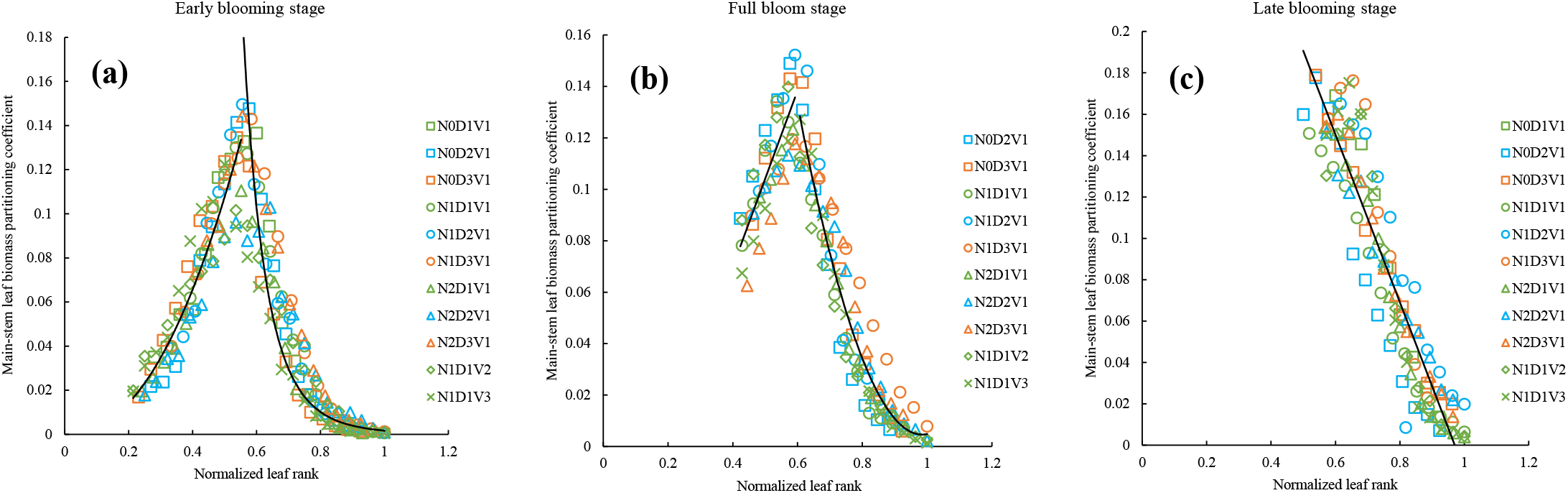
The changes in main-stem leaf biomass partitioning coefficient with normalized leaf rank in the early blooming stage (a) and in the full bloom stage (b) and in the late blooming stage (c) under different treatments.

The main-stem leaf biomass partitioning coefficient of rapeseed in early blooming stage under different treatments showed a trend of increasing first and then decreasing (Fig. 5). The values of *PC*_*EB*_(*LR*) of different treatments with normalized leaf rank were fitted by a power function and a quadratic function (Equation 12 and Fig. 5A). The values of *PC*_*FB*_(*LR*) of different treatments with normalized leaf rank were fitted by a linear function and a quadratic function (Equation 13 and Fig. 5B). The values of *PC*_*LB*_(*LR*) of different treatments with normalized leaf rank were fitted by the quadratic function (Equation 14 and Fig. 5C).

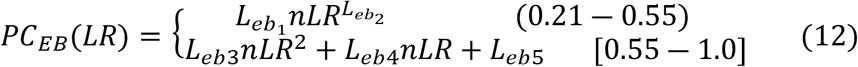

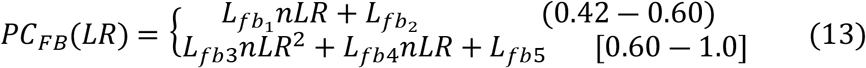

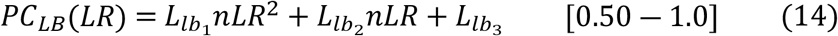

where *PC*_*EB*_(*LR*), *PC*_*FB*_(*LR*) and *PC*_*LB*_(*LR*) are the leaf biomass partitioning coefficient at different leaf rank during the early blooming stage, full blooming stage, and late blooming stage, respectively (g g^−1^); *nLR* is the normalized leaf rank; 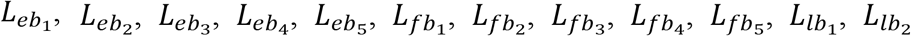 and 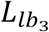 are model parameters. Detailed parameter estimation and significance test are shown in Table 3.

##### 3.2.2.4.Leaf rank boundaries in mature stage: 0.71-1.0

During the mature stage, the biomass of rapeseed plants is primarily transferred to the siliques, with most of the main stem leaves initiating senescence and abscission (Fig. 1A). Consequently, there is a significant reduction in leaf numbers, while the leaf biomass partitioning coefficient reaches its maximum values throughout the entire growth cycle (Fig. 1B and Fig. 6). The leaf rank interval in the mature stage was (0.71-1.0].

**Figure 6.**
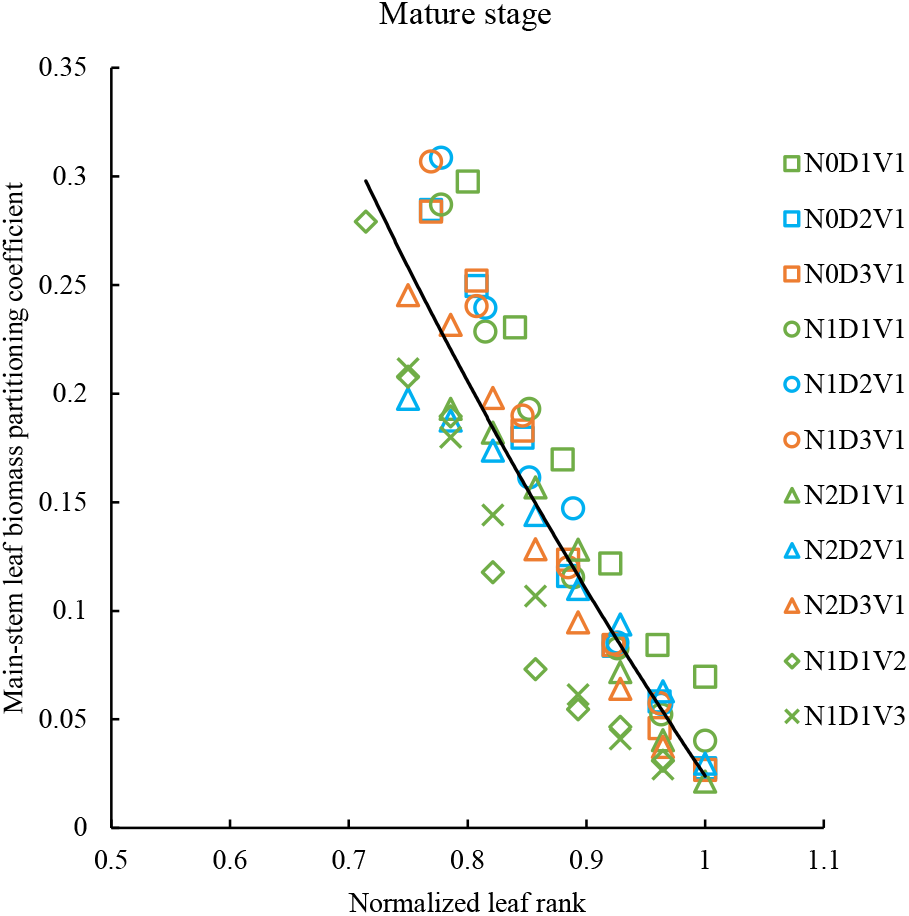
The changes in main-stem leaf biomass partitioning coefficient with normalized leaf rank in the mature stage

The main-stem leaf biomass partitioning coefficient of rapeseed in mature stage under different treatments all showed decreasing trend (Fig. 6). The values of *PC*_*M*_ (*LR*) of different treatments with normalized leaf rank were fitted by the logarithmic function (Equation 15 and Fig. 6).

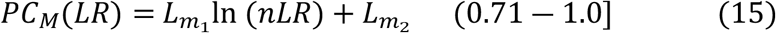

where *PC*_*M*_(*LR*) is the leaf biomass partitioning coefficient at the different leaf rank during the mature stage of rapeseed (g g^−1^); *nLR* is the normalized leaf rank; 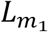 and 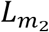 are model parameters. Detailed parameter estimation and significance test are shown in Table 3.

### 3.3 Model Validation

The model developed in this paper are validated using independent experimental data in the 2012-2013 and 2014-2015, and the results showed that the correlation (*r*) of simulation and observation values for the average leaf biomass partitioning coefficient and the leaf biomass partitioning coefficient under different treatments during different growth stage all had significant level at *p*<0.001. The absolute values of the average absolute difference (*d*_*a*_) are all less than 0.011 g g^−1^, the values of the root mean square error (*RMSE*) are all less than 0.032 g g^−1^, the ratio of *d*_*a*_ to the average observation (*d*_*ap*_) are all less than 11.312%, which indicated that the observed and simulated leaf biomass partitioning coefficient were all close to the 1:1 line. The average absolute difference (*d*_*a*_) values for average leaf biomass partitioning coefficient (*PC*_*AL*_), leaf biomass partitioning coefficient at seedling stage (*PCs*), and at budding and bolting stage (*PC*_*Bb*_), and at blooming stage (*PC*_*B*_), and at mature stage (*PC*_*M*_) under different treatments are −0.009 g g^−1^, −0.010 g g^−1^, 0.007 g g^−1^, 0.002 g g^−1^, and −0.011 g g^−1^, respectively. The root mean square error (*RMSE*) values for *PC*_*AL*_, *PCs, PC*_*Bb*_, *PC*_*B*_, *PC*_*M*_ are 0.031 g g^−1^, 0.032 g g^−1^, 0.016 g g^−1^, 0.017 g g^−1^, and 0.032 g g^−1^, respectively. The ratio of *d*_*a*_ to the average observation (*d*_*ap*_) of simulation and observation values for *PC*_*B*_ is less than 5 %, the *PC*_*AL*_ and *PC*_*M*_ is less than 10 %, the PCs and *PC*_*Bb*_ are less than 12 %, which indicate that the models constructed in this study have a good performance and reliability (Fig. 7 and Table 4).

**Table 4.** Comparison of statistical parameters of observation and simulation in main-stem leaf biomass partitioning coefficient models in 2012-2013 and 2014-2015.

| Architectural parameter | Statistic parameters of simulation and observation |  |  |  |  |
| --- | --- | --- | --- | --- | --- |
| | $n$ | $d_a$ (g g<sup>-1</sup>) | $d_{ap}$ | $RMSE$ | $r$ |

|  |  | <sup>1)</sup> | (%) | (g g<sup>-1</sup>) |  |
| --- | --- | --- | --- | --- | --- |
| Normalized leaf rank (nLR) | 1859 | -0.08 | 13.559 | 0.200 | 0.657*** |
| Leaf biomass partitioning coefficient<br>at seedling stage (PCs) | 291 | -0.010 | 11.236 | 0.032 | 0.943*** |
| Leaf biomass partitioning coefficient<br>at budding and bolting stage (PC<sub>Bb</sub>) | 381 | 0.007 | 11.312 | 0.016 | 0.936*** |
| Leaf biomass partitioning coefficient<br>at blooming stage (PC<sub>B</sub>) | 775 | 0.002 | 3.077 | 0.017 | 0.936*** |
| Leaf biomass partitioning coefficient<br>at mature stage (PC<sub>M</sub>) | 84 | -0.011 | 8.661 | 0.032 | 0.938*** |

**Figure 7.**
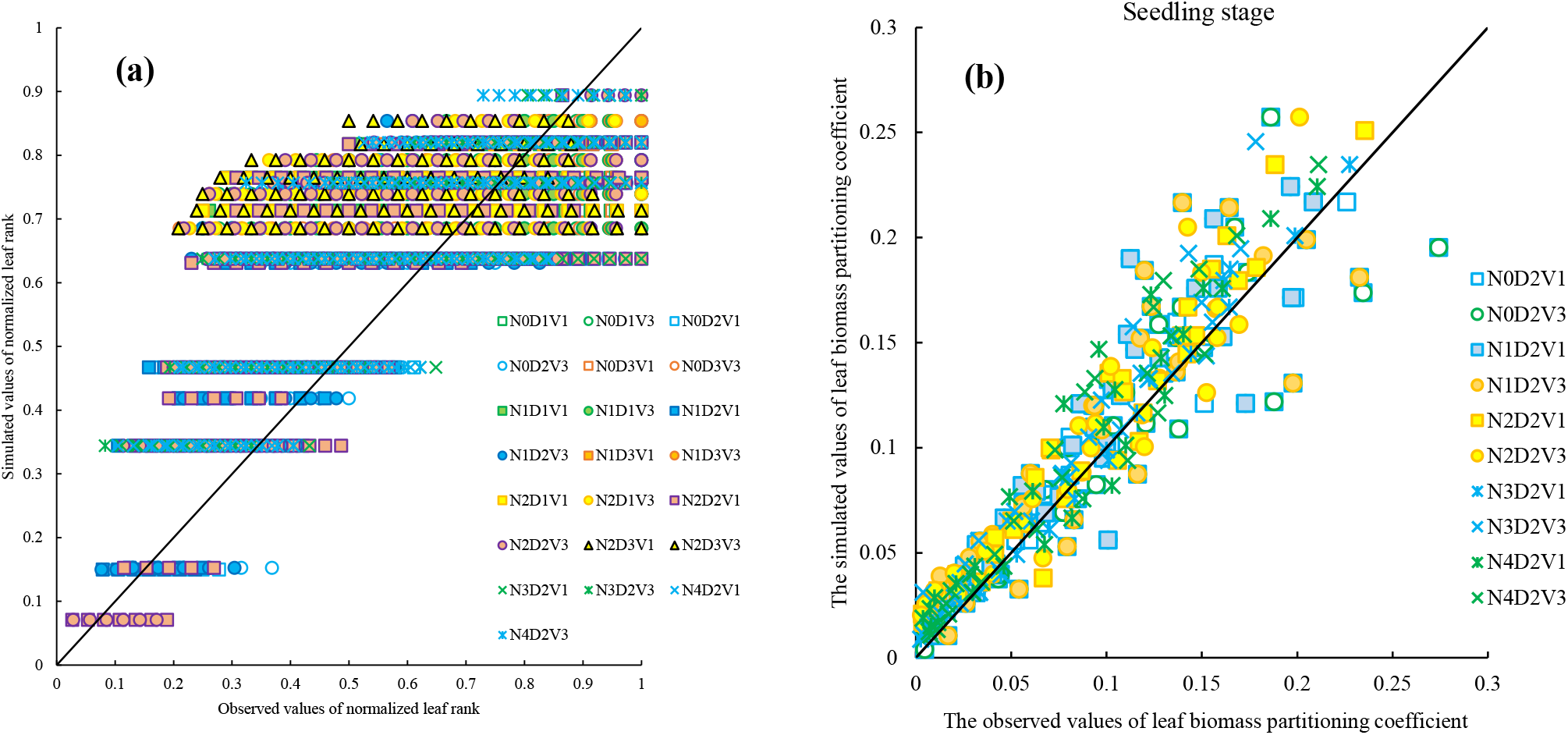
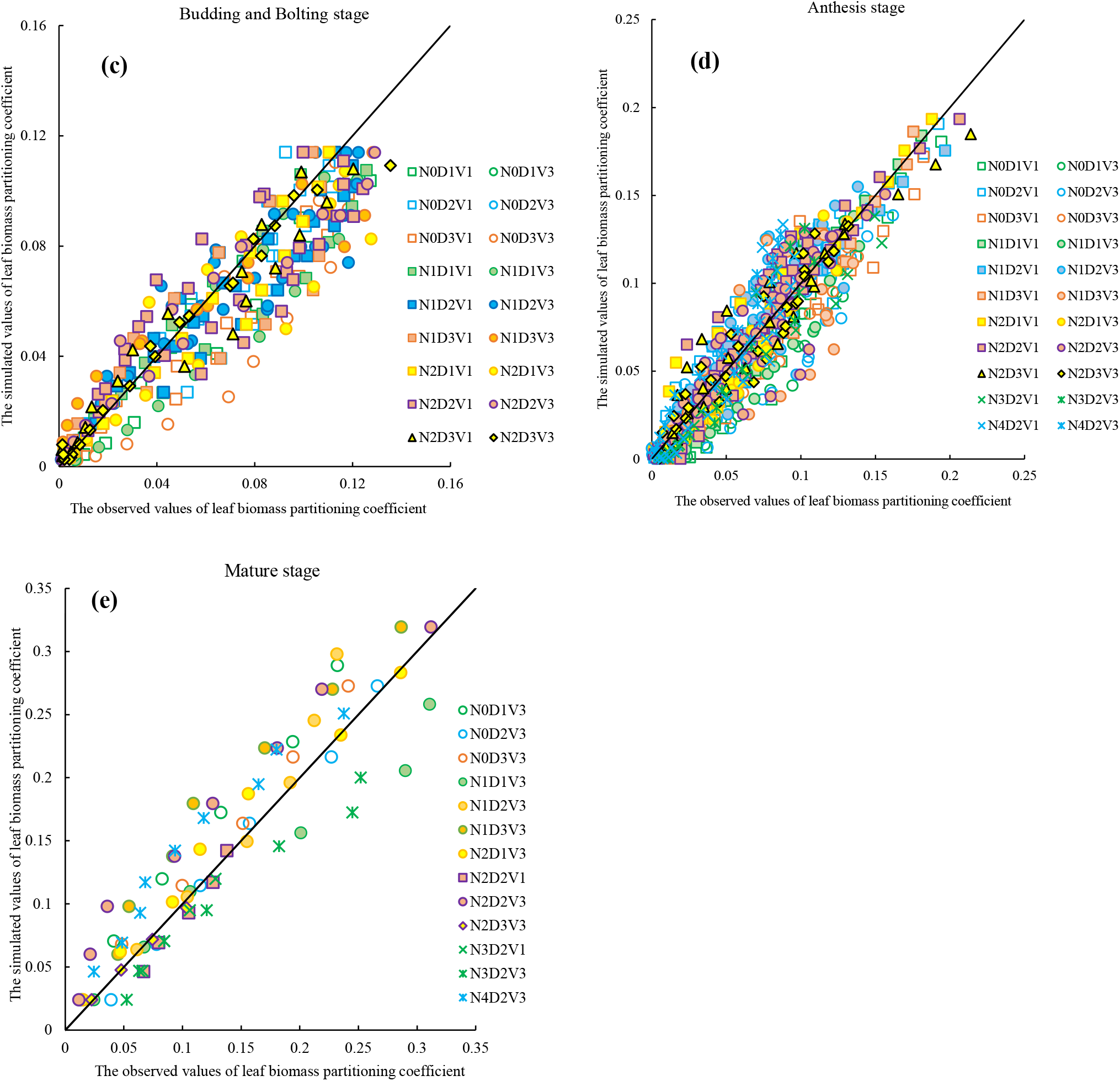
Comparison of the main-stem average leaf biomass partitioning coefficient (a) and the leaf biomass partitioning coefficient observed values with the simulated values at (b) seedling stage and (c) budding and bolting stage and (d) anthesis stage and (e) mature stage under different treatments in 2012-2013 and 2014-2015.

## 4. Discussion

### 4.1 Effects of nitrogen fertilization on leaf biomass partitioning in rapeseed

The vegetative phase in rapeseed establishes the photosynthetic infrastructure critical for yield formation, with leaf biomass accumulation serving as both a driver and indicator of reproductive potential (Zhang *et al*., 2020). Nitrogen (N) fertilization significantly enhances photosynthetic performance by increasing ribulose-1,5-bisphosphate carboxylase/oxygenase (RuBisCO) activity and leaf area index (LAI), thereby improving canopy light interception efficiency (Poorter and Nagel, 2000; Zhang *et al*., 2016). Conversely, N-deficient conditions induce leaf structural constraints, including reduced mesophyll cell density and impaired stomatal conductance constrain light harvesting and carbon fixation, creating metabolic bottlenecks during early developmental stages (Kapen *et al*., 1998; Leng *et al*., 2000). During reproductive transition, leaves undergo functional specialization from autonomous carbon sources to transient sinks, a shift modulated by silique-dominated assimilate competition (Rao *et al*., 1991; Boonman *et al*., 2006; Brouwer *et al*., 2012; Brunel-Muguet *et al*., 2013). This sink hierarchy precipitates architectural trade-offs, as developing reproductive structures shade lower leaves, reducing their photosynthetic active radiation (PAR) interception and accelerating senescence through cytokinin redistribution (Grbićand Bleecker, 1995; Gabrielle *et al*., 1998; Boonman *et al*., 2006; Brouwer *et al*., 2012). This architectural trade-off is genotype-dependent, with cultivars exhibiting differential strategies in maintaining source-sink balance through flowering progression (Dai *et al*., 2001; Hu *et al*., 2002). Leaf size and plant numbers during this phase serve as key predictors of subsequent yield components.

Adequate nitrogen supply plays a crucial role in shaping biomass allocation patterns within rapeseed plants, particularly in their leaves. Optimal nitrogen levels promote growth and development, resulting in increased leaf biomass. This increase in leaf biomass is often accompanied by changes in biomass allocation, with a higher proportion of total biomass being allocated to leaves compared to other plant organs (Malagoli *et al*., 2005). Moreover, the application of nitrogen fertilizer can influence the structural and physiological characteristics of rapeseed leaves. It can lead to larger leaf area, higher chlorophyll content, and improved photosynthetic activity. These alterations in leaf morphology and physiology contribute to enhanced carbon assimilation and ultimately higher biomass production in rapeseed plants (Liu *et al*., 2020). However, it is important to note that excessive nitrogen application can have negative effects on biomass allocation in rapeseed leaves. It can disrupt nutrient uptake balance, heightened susceptibility to diseases, and environmental stress. Furthermore, nitrogen oversupply can promote vegetative growth at the expense of reproductive development, potentially compromising seed yield and quality (Gaju *et al*., 2014).

### 4.2 Effects of plant density on leaf biomass partitioning in rapeseed

Planting density represents a critical agronomic factor impacting crop productivity and resource use efficiency, with significant implications for leaf biomass partitioning patterns in rapeseed (Leach *et al*., 1999; Momoh and Zhou, 2001). Studies have indicated that rapeseed plants cultivated at higher densities generally exhibit accelerated growth rates and enhanced biomass accumulation compared to those at lower densities (Li, 2015). Nevertheless, the complex mechanisms through which transplanting density modulates biomass partitioning in rapeseed leaves necessitate more in-depth exploration.

Elevated planting densities typically intensify inter-plant competition for limited resources such as light, water, and nutrients. As an adaptive strategy, rapeseed plants often modify their biomass allocation patterns, particularly in leaf organs, to cope with intensified competitive pressures. For instance, Kuai *et al*. (2022) revealed that increasing planting density induced notable shifts in biomass partitioning, where lower densities promoted a higher proportion of biomass allocated to leaves. Concurrently, higher-density plantings were associated with the development of taller and more elongated plant architectures, accompanied by increased Leaf Area Index (LAI) and Radiation Use Efficiency (RUE) across successive growth stages (Kuai *et al*., 2016). These observations indicate that planting density not only regulates biomass distribution but also influences plant morphological traits and resource utilization efficiency.

Understanding how planting density affects rapeseed leaf growth is crucial for elucidating the implications for biomass accumulation and distribution along the main stem (Leng *et al*., 2004). Experimental evidence demonstrates that varying planting densities result in distinct patterns of leaf biomass partitioning: higher densities tend to reduce biomass allocation to lower leaf ranks while promoting greater accumulation in upper leaf positions (Brunel-Muguet *et al*., 2013). In contrast, lower planting densities facilitate more equitable biomass distribution along the main stem, likely due to diminished inter-plant competition for essential resources. During the vegetative growth stage, leaves serve as primary organs for biomass accumulation, playing a pivotal role in establishing the foundation for subsequent grain formation (Liao and Guan, 2002; Gregersen, *et al*., 2008). Adequate biomass accumulation during this stage is therefore critical for achieving optimal grain yield in later developmental stages.

### 4.3 Mechanistic modelling of biomass accumulation and distribution in rapeseed

The quantitative prediction of biomass allocation patterns has emerged as a critical frontier in rapeseed yield, with researchers worldwide employing diverse modelling approaches to dissect these processes. Notable models include the DAR95 (DAISY-Rape) model developed by Petersen *et al*. (1995), which utilizes the distribution coefficient method to illustrate biomass allocation to different plant organs, including seeds. Additionally, Liao *et al*. (2000) employed logistic curve analysis to investigate biomass allocation in winter rapeseed, highlighting the preferential allocation to leaves before winter and siliques during seed development. Liu *et al*. (2005) and Wang *et al*. (2008) further advanced this field by introducing dynamic simulation models that link aboveground allocation indices to physiological developmental time, offering temporal insights into biomass redistribution.

Tang *et al*. (2007) explored cultivar- and nitrogen fertilizer-driven variations in biomass distribution, though their model’s generality could be enhanced by integrating additional environmental variables such as temperature and light intensity. Building on these foundations, Cao *et al*. (2006) developed a mechanistic model incorporating photosynthetic processes and microclimatic factors to simulate rapeseed growth dynamics. Robertson *et al*. (2016) applied APSIM platform to create APSIM-Canola, a comprehensive model detailing phenological and morphological development. Conversely, the EPR95 (EPIC-Rape) model by Kiniry *et al*. (1995) offers broad applicability across crops and environments but relies on effective accumulated temperature, a limitation that may compromise its accuracy in predicting rapeseed development under heterogeneous climatic conditions despite its user-friendly interface. Current challenges in biomass modelling necessitate the development of more integrative frameworks that capture the dynamic interplay between genetic, physiological, and environmental drivers of biomass partitioning. Future research should prioritize mechanistic models capable of resolving how planting density, nutrient availability, and climatic variables modulate allocation patterns across vegetative (e.g., leaves, stems) and reproductive organs. For instance, refining leaf biomass partitioning coefficient models, such as those developed herein, can enhance understanding of vertical biomass distribution along the main stem, a critical factor in optimizing radiation interception and resource use efficiency.

The present study builds on prior characterizations of biomass allocation across organs (e.g., main stem, total leaves, and branches) by introducing a hierarchical model of leaf rank-specific biomass partitioning. This approach addresses a key gap in existing literature by dissociating biomass allocation dynamics among upper and lower leaf ranks, which are differentially influenced by light competition and source-sink relationships. In this study, the validation results confirms the model reliably simulates leaf biomass partitioning across key developmental stages. While the model demonstrates robust overall performance, elevated *d*_*ap*_ values (>10%) occur specifically during seedling establishment (PCs), stem elongation (PC_Bb_), and across normalized leaf ranks (nLR). We attribute this phenomenon to two interrelated factors characteristic of rapeseed’s vegetative growth phase: First, inherently low biomass partitioning coefficients during rapid leaf expansion amplify the relative impact of minor absolute errors. Second, subtle environmental variations (such as microclimate gradients or resource availability), induce substantial biological variability in foliar development patterns, particularly at individual leaf positions. These combined effects naturally increase dispersion between observed and simulated values during these dynamic growth stages.

By integrating empirical data with process-based algorithms, these models not only advance theoretical understanding of rapeseed functional ecology but also provide actionable tools for precision agriculture, enabling farmers to optimize planting density and nitrogen management to enhance yield potential. Prospective validation under diverse field conditions will be essential to refining model parameters and ensuring their scalability. Ultimately, such advancements in functional-structural modelling will strengthen our capacity to predict crop performance under climate change scenarios, fostering the development of resilient and resource-efficient rapeseed production systems.

## 5. Conclusions

This paper presented a leaf biomass partitioning coefficient model of main-stem at different leaf ranks in rapeseed, through analyzing three years of field experimental data with three cultivars, five fertilizer levels, and three planting density levels. We established a model for the leaf biomass partitioning coefficient of the main-stem at different growth stage by analyzing the quantitative relationship between the leaf biomass partitioning coefficient and the normalized leaf rank (*nLR*). The validation of descriptive models using independent experimental data demonstrated robust predictive accuracy, with statistically significant correlations (*r* > 0.9, *p* < 0.001) between simulated and observed values, exhibiting a mean absolute difference (*d*_*a*_) below 0.011 g g^−1^ and stage-specific normalized error ratios (*d*_*ap*_) ranging from 3.077% (mature stage) to 13.599% (seedling stage), complemented by *RMSE* values consistently under 0.032 g g^−1^, collectively affirming the model’s reliability in simulating main-stem leaf biomass partitioning coefficients across hierarchical leaf ranks in rapeseed.

## Acknowledgements

This work was jointly supported by the grants from the National Natural Science Foundation of China (32201664, 31871522, 31601223, and 31471415), the Fund of Jiangsu Academy of Agricultural Sciences (6111648), and the Jiangsu Provincial Key R&D Program (BE2023302). We sincerely thank Xiufang Wei, Yanbin Yue, Kunya Fu, Chunhuan Feng, Weitao Chen, Chuwei Song, Sijun Ge, Qian Zhang, Qian Wan, and Shujiang Feng for their technical assistance in the field.

## Appendix A.

Physiological day of development (*PDD*) which was used to drive the model (Gao *et al*., 1992), defined as follows

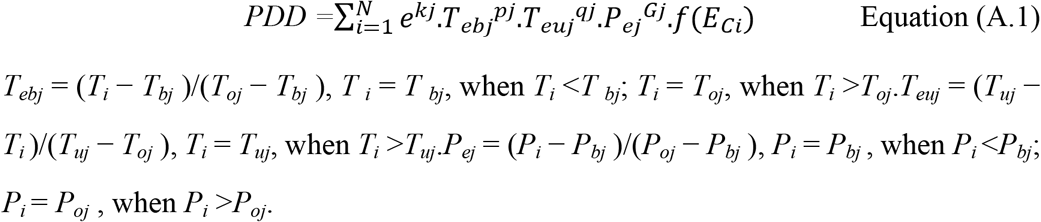

where the *N* is the number of days covering the stage, the *k*_*j*_ is basic development parameter which is determined by cultivar heredity, the *T*_*ebj*_ and the *T*_*euj*_ are the effective factors for lower and upper temperature, respectively, the *pj* and the *q*_*j*_ are the genotypic coefficient of temperature effects, the *P*_*ej*_ is the effective factor of photoperiod, *G*_*j*_ is the genotypic coefficient of photoperiod effects, and f (*E*_*Ci*_) is the effective function of agronomic practice factors for rapeseed. *T*_*i*_ is the daily mean temperature (°C) in the *j*th stage, *T*_*bj*_, *T*_*oj*_ and *T*_*uj*_ are lower, optimum, and upper limit temperature (°C) demanded in the *j*th stage for rapeseed, respectively, and *P*_*bj*_, *P*_*oj*_ are the critical and optimum day length (h) demanded in *j*th stage for rapeseed, respectively. *PDD* 0, 1, 2, 3, and 4 represent sowing, seedling, bolting, flowering, and maturity, respectively.

## Notes

### Competing Interest Statement

The authors have declared no competing interest.

